# Soquelitinib, A Selective Inhibitor of Interleukin-2-Inducible T Cell Kinase (ITK), is Active in Several Murine Models of T Cell-Mediated Inflammatory Disease

**DOI:** 10.1101/2023.10.27.564296

**Authors:** Lih-Yun Hsu, James T. Rosenbaum, Sarah Lindner, Yannick Allanore, Anne Cauvet, Rahul D. Pawar, Dan Li, Marcel Van Den Brink, Richard A. Miller

## Abstract

Interleukin-2-inducible T cell kinase (or ITK) is a tyrosine kinase predominantly expressed by T lymphocytes. It plays a major role in T cell activation, differentiation, and receptor signaling. Studies in mouse models have established that the absence or inhibition of ITK reduces the secretion of Th2 and Th17 cytokines, while favoring the production of Th1 cytokines and the differentiation of T cells to regulatory T cells (T regs). We recently characterized the activity of soquelitinib (SQL), a selective, covalent inhibitor of ITK *in vitro* using mouse or human T cells and *in vivo* in murine models of cancer. We hypothesized that selective pharmacologic blockade of ITK could attenuate a variety of T cell-mediated inflammatory diseases including murine models that resemble asthma, pulmonary fibrosis, systemic sclerosis, psoriasis, and acute graft versus host disease. In each model, SQL demonstrated substantial ability to ameliorate the disease process along with effects on cytokines and/or T cell subsets consistent with the reported function of ITK. Our studies demonstrate that selective inhibition of ITK by SQL is potentially a novel therapeutic approach for several forms of T cell-mediated, inflammatory diseases.

## INTRODUCTION

The T cell receptor (TCR) plays an essential role in T cell function. Once activated, the TCR signals through a chain of transmembrane and intracellular molecules including interleukin-2-inducible T cell kinase or ITK which is a member of the TEC family of tyrosine kinases^1–5^. In addition to T cells, ITK has limited expression including in some mast cells, some innate lymphoid cells (ILCs)^6^ and NK cells. The critical function of ITK has been elucidated in part by studies in knockout or gene altered mice^7,8^. These observations indicate that inactivation or deletion of ITK reduces Th2 and Th17 cells and related cytokines without marked inhibition of Th1 cells. These findings presumably result from unimpeded activity of RLK (resting lymphocyte kinase) which plays a redundant function in Th1 cell development.

We recently reported the characterization of soquelitinib (SQL), a highly selective, covalent inhibitor of ITK^9^. We observed that SQL is active in several mouse models of cancer, including malignancies that do not express ITK. SQL treatment results in activation of cytolytic CD8 cells in part via Th1 cells which produce interferonγ (IFNγ). In addition, SQL reduces the expression of several markers of T cell exhaustion such that the T cell-mediated, anti-cancer immune response is potentiated.

The properties of SQL make it potentially useful in the management of several human immune-mediated diseases which include atopic diseases like asthma and atopic dermatitis which are mediated by Th2 cytokines; fibrosis in which Th2 cells have been strongly implicated^10,11^; psoriasis in which Th17 plays a major role^12^; and acute graft versus host disease (aGVHD) in which the T cell receptor, Th17^13^, and regulatory T^14^ cells (Tregs) contribute. Accordingly, we studied mouse models that resemble these human diseases. We report that SQL is effective in murine models of asthma, pulmonary fibrosis, systemic sclerosis, psoriasis, and aGVHD by mechanisms consistent with the biologic effects described above.

## RESULTS

### Soquelitinib is efficacious in murine models of asthma

ITK has been considered an attractive target for modulating Th2-driven inflammatory diseases such as allergic asthma due to its pivotal role in mediating the production of Th2-associated cytokines. ITK-deficient mice display an impairment in mounting effective Th2-mediated immunity, and thereby are protected from allergic asthma^7,8^. To assess whether SQL is effective in a Th2-driven disease model, we investigated an ovalbumin (OVA)-induced mouse acute asthma model^15^ (Figure 1A) in which CD4+ Th2 cells represent the predominant cell infiltrate in the lung tissue. Dexamethasone given intraperitoneally (IP) at a dose of 1 mg/kg was used as a positive control in these studies.

**Figure 1.**
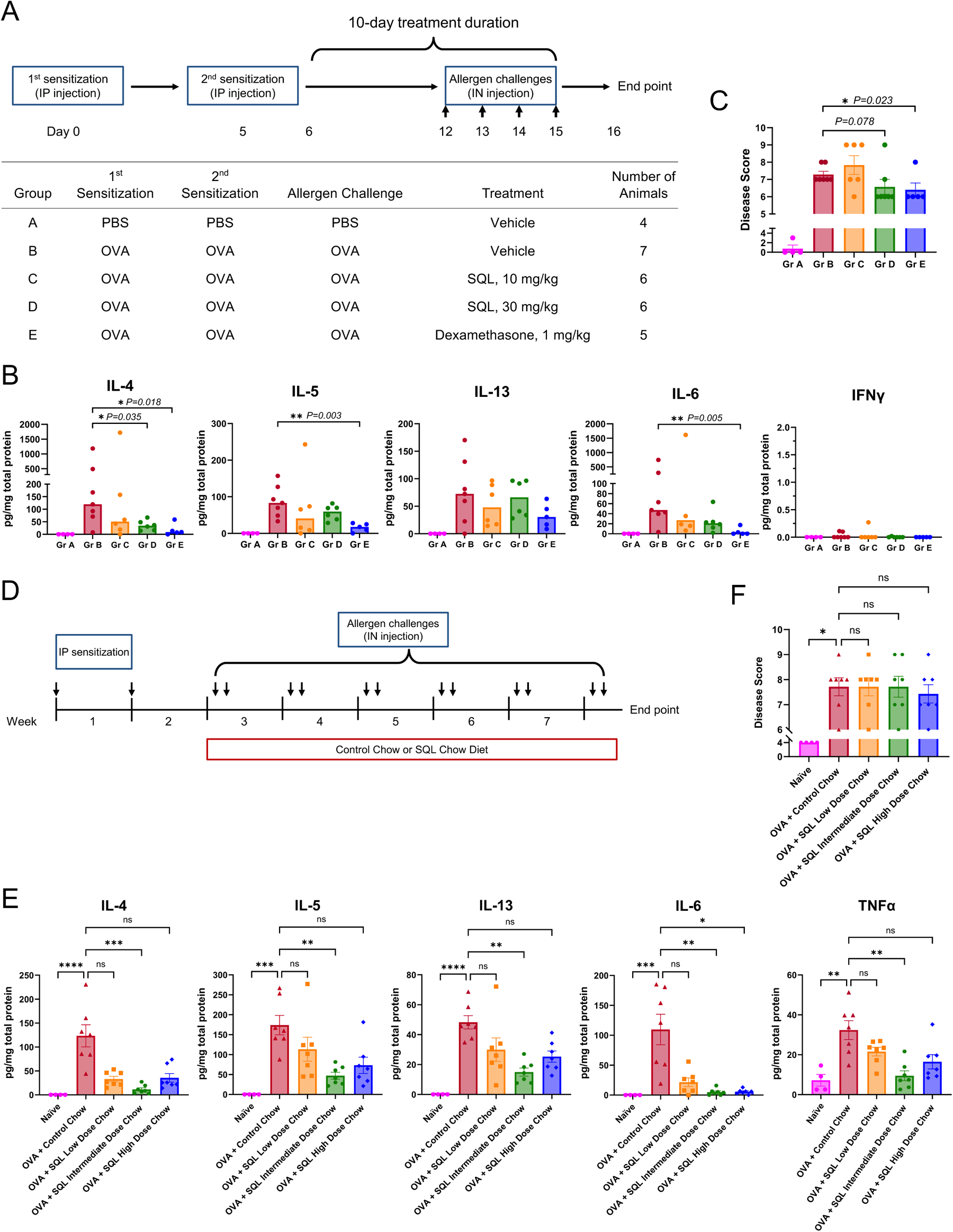
Soquelitinib attenuates OVA-induced airway inflammation. (A & D) Schematics illustrate experimental design of OVA-induced acute asthma (A) and chronic asthma (D) in mice. (B & E) Inhibition of Th2-associated and inflammatory cytokines by soquelitinib in acute asthma (B) and chronic asthma models (E). The levels of indicated cytokines in BALF were measured by multiplex MSD. (C & F) Total disease scores for assessment of lung injury are shown. (E) Intermediate dose of soquelitinib chow diet exhibits the strongest inhibition of Th2-associated and inflammatory cytokines. Statistical significance was determined by Mann-Whitney U test. Data (mean±SEM) are representative of two independent experiments with 4 to 7 mice per group. Each dot in a bar graph represents one mouse. IP=intraperitoneal; IN=intranasal. * p<0.05; ** p<0.01; *** p<0.001; **** p<0.0001.

Consistent with the previous reports^15^, we demonstrated that repeated intranasal OVA challenges induced elevated levels of Th2-associated cytokines such as IL-4, IL-5, and IL-13, but not the Th1-associated cytokine, IFNγ, in the broncho-alveolar lavage fluid (BALF) samples from group B mice (Figure 1B). On day 6, mice were treated by oral gavage with either vehicle alone, or one of two different doses of SQL, 10 mg/kg or 30 mg/kg. Doses were administered once daily for ten consecutive days in a volume of 200μl. SQL treatment resulted in a trend toward reduced levels of IL-4, IL-5, and IL-13 in the BALF samples, as compared to the vehicle-treated group (Figure 1B). The decrease in IL-4, a central driver of Th2-driven inflammation, reached statistical significance in mice receiving intermediate dose of SQL at a dose of 30 mg/kg by oral gavage once daily.

Within the limits of accuracy of the assay, SQL treatment did not result in skewed cytokine production to Th1-associated IFNγ, as IFNγ remained barely detectable in both groups C and D. We also observed that SQL-treated groups showed moderate reduction of a broad inflammatory cytokine, IL-6. As shown in Figure 1C, histopathological disease was reduced by dexamethasone and it tended to be reduced by the 30 mg/kg dose of SQL (p=0.078) despite one outlier and a small sample size. Representative histology is shown in supplemental Figure 1A.

To expand the findings, we evaluated the effect of varying concentrations of SQL in a chronic mouse asthma model^16^ where OVA allergen was administrated intranasally multiple times over a period of 5 1/2 weeks (Figure 1D). During the same period, OVA-sensitized mice were treated with either control chow or one of three different doses (see Methods) of SQL-formulated chows. As in acute asthma, the model is associated with an increase in Th2 cytokines (IL-4, 5, and 13) detectable in BALF. All 3 doses of SQL reduced the level of these cytokines (Figure 1E). Two other cytokines, IL-6 and TNFα, were also reduced by SQL (Figure 1E). None of the treatments had a statistically significant effect on the histologic grading in this model (Figure 1F). Together, we have demonstrated that SQL treatment in either acute or chronic asthma settings can suppress the production of several Th2 type cytokines and inflammatory cytokines.

### Soquelitinib reduced bleomycin-induced fibrosis

The effect of SQL on pulmonary fibrosis was studied in a murine, bleomycin-induced model of pulmonary fibrosis. Five groups of mice were studied as shown in Figure 2A. All groups were challenged with bleomycin delivered via the oro-pharynx on days 0 and 6 except for the Group 1 mice that received only phosphate buffered saline (PBS). Group 2 received treatment with just the vehicle diluent. Groups 3 and 4 were treated with one of two doses of SQL on days 7 through 21 at either 30 mg/kg or 10 mg/kg given by gavage orally once per day. Group 5 received nintedanib, an FDA-approved tyrosine kinase inhibitor for treatment of idiopathic pulmonary fibrosis^17^. Nintedanib inhibits receptors for vascular endothelial growth factor, fibroblast growth factor and platelet derived growth factor. Lung weights were increased by bleomycin challenge (Figure 2B). Both doses of SQL or nintedanib limited this increase. Additionally, Figures 2C and 2D show that the lower dose of SQL or nintedanib reduce the leukocytic infiltrate in the BALF, and all 3 treatments reduced the Ashcroft score^18^, which is indicative of structural lung damage, compared to vehicle treated mice. Representative hematoxylin and eosin histology is shown in supplemental Figure 2A. Fibrosis was reduced by either dose of SQL, more consistently than the benefit from nintedanib (Figure 2E). Representative trichrome staining for fibrosis is shown in supplemental Figure 2B. The caval lobe was used to investigate gene expression in the injured lung using qPCR. As shown in Figures 2F and 2G, the major transcription factor controlling the expression of Th2 cytokines, GATA-3, is reduced by the 30 mg/kg dose of SQL, and two mRNAs associated with fibrosis, MMP2 (matrix metalloproteinase 2) and TGFβ (transforming growth factor beta), are reduced by the 30 mg/kg dose of SQL. Together, our findings indicated that SQL treatment enabled downregulation of genes that exert important functions in pulmonary fibrosis.

**Figure 2.**
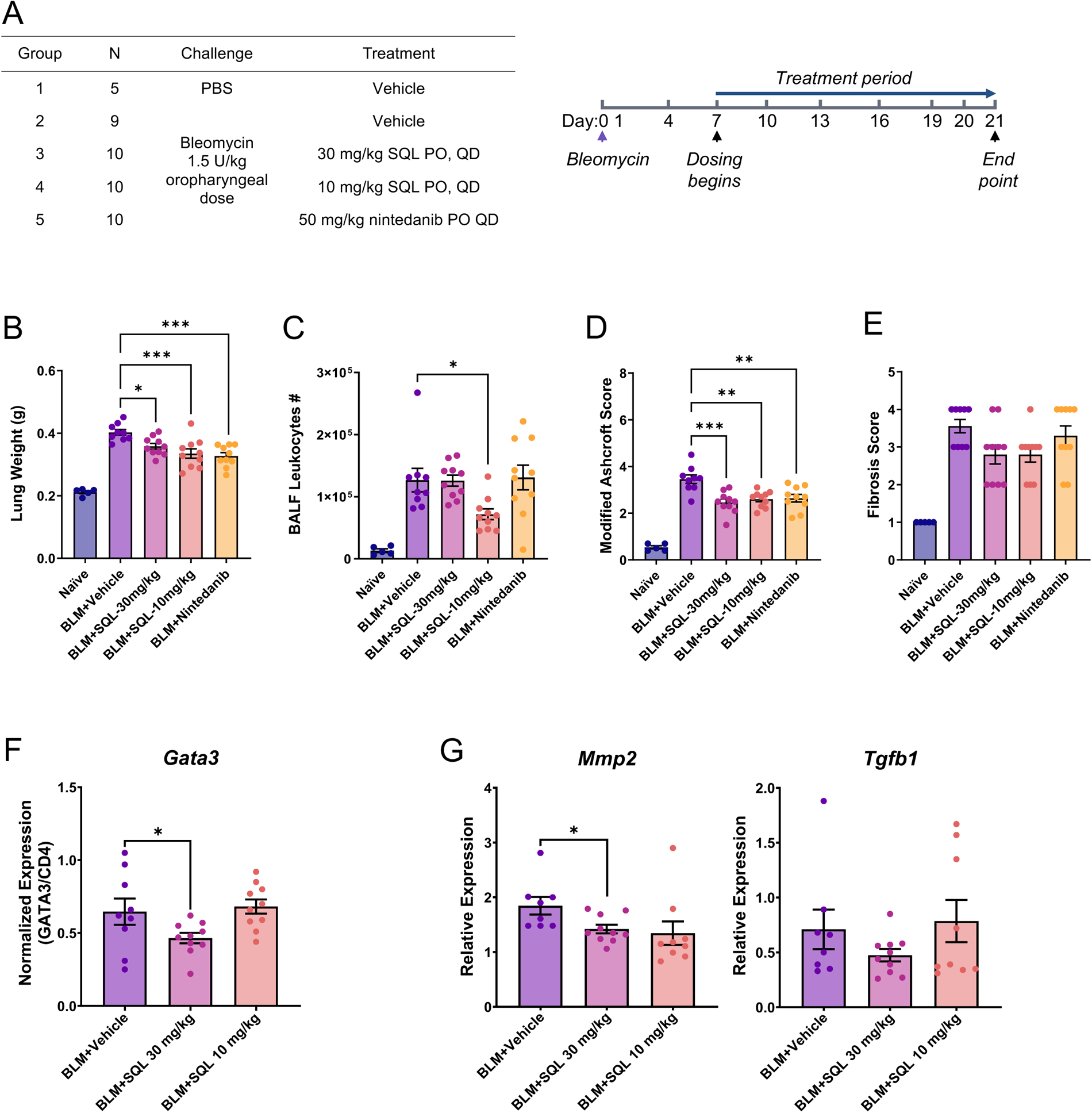
Soquelitinib treatment protects mice against bleomycin(BLM)-induced lung fibrosis. (A) Table of the experimental groups and schematic of the study. (B) Absolute lung weight among different groups is shown. n=4–10 per group. Statistical significance was determined by one-way ANOVA. (C) Soquelitinib reduces recruitment of leukocytes into the lung. Infiltrating leukocyte number collected from BALF samples is shown. (D) Soquelitinib significantly decreases lung structural damage. Modified Ashcroft scores for the severity of fibrosis are shown. Statistical significance was determined by one-way ANOVA. (E) Soquelitinib treatment results in a moderate decrease in lung fibrosis. Fibrosis scores quantified by evaluating the Masson’s Trichrome (MT)-stained sections are shown. (F) Expression of the Th2 lineage defining transcription factor, GATA-3 and (G) two fibrosis-associated transcripts (MMP2 and TGFβ1) in lung tissues were determined by quantitative RT-PCR from 9–10 mice in each group. CD4 was used for normalization of infiltrating CD4 T cell numbers (F). Fold changes of two fibrosis-associated genes relative to PBS control are shown (G). Each dot in a bar graph represents one mouse. * p<0.05; ** p<0.01; *** p<0.001.

### Soquelitinib is effective in a mouse model of systemic sclerosis

The effect of SQL in a murine model of systemic sclerosis was studied in mice that over express the transcription factor, Fra-2 (*FosL2*, an AP1 transcription factor). This mutation results in widespread inflammation, vasculopathy, pulmonary fibrosis, and pulmonary hypertension. The mice develop vascular injury which precedes fibrosis as is seen in systemic sclerosis^19,20^. The model and techniques for its assessment have been previously described^19,20^. In the description of results, we refer to these mice that over express Fra-2 simply as Fra-2 mice. When the mice were 11 weeks old (before the onset of clinical disease) animals were divided into four groups for treatment: wild type controls (n=6), Fra-2 mice receiving regular chow (n=15), Fra-2 mice receiving low dose SQL-formulated chow (n=14), and Fra-2 mice receiving the intermediate dose SQL-formulated chow (n=12). Treatment continued for 7 weeks.

Fra-2 mice are known to lose weight as the disease progresses. As shown in Figure 3A, the mice receiving the intermediate dose of SQL maintained weight equal to healthy controls, in contrast to Fra-2 mice in the control group or Fra-2 mice receiving the lower dose of SQL chow. None of the healthy control mice died. There were four deaths in the Fra-2 mice eating regular chow, 3 deaths in the Fra-2 mice receiving the low dose SQL chow, and 2 deaths in the Fra-2 group receiving the intermediate dose of SQL chow (see Kaplan-Meier curve, Figure 3B).

**Figure 3.**
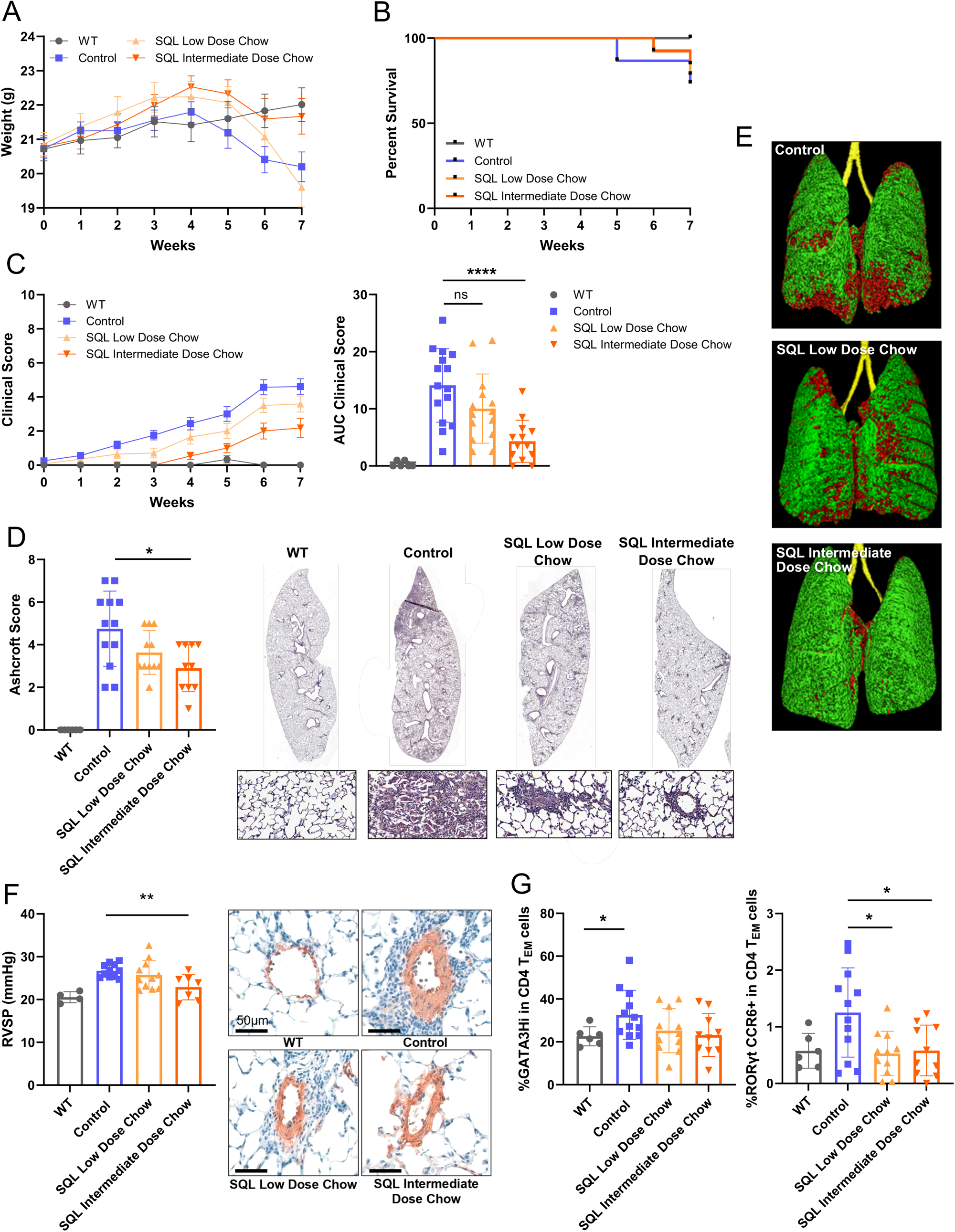
Soquelitinib reduces disease in the Fra-2 model of systemic sclerosis. (A) Wild type mice and mice receiving the higher dose of SQL have better weight gain than the group treated with the lower dose of SQL or the vehicle treated group. (B) Kaplan-Meier survival curve showing slight benefit for the SQL higher dose treated mice. The two groups treated with SQL show results that overlap and thus the two curves are difficult to visually separate. (C) The clinical score is improved by SQL treatment compared to vehicle treated controls. The differences between the high dose treated and the vehicle treated mice become statistically significantly different by two weeks of therapy. The improvement for the high dose treated mice as determined by the area under the curve (AUC) is highly significant (p<0.001). (D) The Ashcroft score is improved by SQL treatment (P<0.05 for those receiving the higher dose of SQL) and as illustrated by the representative histology. (E) Representative micro-CT images of the lungs show benefit from SQL treatment. (F) The right ventricular systolic pressure (RVSP) is lowered by SQL treatment at the intermediate dose (p<0.01). Remodeling of the vessel was assessed by staining for smooth muscle actin and shows an increase in wall thickness associated with perivascular inflammatory infiltrates in the Fra-2 control mice and its reduction by SQL treatment. (G) Expression of GATA-3, the major Th2 transcription factor, is increased in Fra-2 mice compared to controls, and reduced by SQL treatment (p>0.05, not significant). Similarly, the major Th17 transcription factor, RORγt, is increased in Fra-2 mice and the expression is reduced by either dose of SQL (p<0.05). Each dot in a bar graph represents one mouse. WT=wild type. * p<0.05; ** p<0.01; *** p<0.001, **** p<0.0001.

Disease severity was assessed using a ten point scale as previously reported^19,20^. One point was given for the presence of blepharitis, alopecia, skin lesions, abnormal fur texture, hepatomegaly, hunched posture, or impaired mobility. Weight loss of 0 to 5% was given one point, while weight loss of 5 to 10% received 2 points, and weight loss of 10 to 15% received 3 points. As shown in Figure 3C, the Fra-2 group treated with SQL was statistically better in terms of the clinical score compared to the untreated Fra-2 group. The onset of improvement is statistically significant as early as two weeks after the onset of treatment. If the comparison of the clinical score is based on the area under the curve, the group receiving the low dose of SQL chow was better than the untreated group, but the results did not reach statistical significance. The Fra-2 mice receiving the intermediate dose of SQL chow were clearly better than the untreated Fra-2 mice (p<0.0001) as judged by the area under the curve for the clinical score.

The degree of fibrosis as indicated by collagen deposition in the lung was measured by quantifying hydroxyproline. This assay showed that the disease was associated with a marked increase in collagen in the Fra-2 mice compared to the controls. Neither dose of SQL reduced the fibrosis as measured by this index. By histopathology, however, the Ashcroft score was significantly better in the Fra-2 mice which received the intermediate dose of SQL chow (p=0.015). Figure 3D illustrates this graphically and includes representative histology. The Ashcroft score is based on scoring 20 representative lung sections for fibrosis on a scale of 1 to 8. This improvement could not be definitively confirmed by micro-CT, although there was a trend in that direction. Figure 3E shows representative CT images indicating the improvement. The right ventricular systolic pressure (RVSP) was measured by right heart catheterization just prior to euthanasia at week 18. As in patients with systemic sclerosis, the mice develop an elevated RVSP due to pulmonary artery hypertension^21^. Again, the mice receiving the intermediate dose of SQL chow were improved compared to untreated controls (p=0.004, Figure 3F). Together, these results demonstrate that SQL reduced pulmonary fibrosis and had a favorable impact on the disease process.

Finally, we used flow cytometry to assess T cell subsets in the spleen of Fra-2 mice. As shown in Figure 3G, and consistent with known effects of either SQL or ITK deletion, SQL treated Fra-2 mice had lower levels of Th2 (p=0.056) and Th17 T cells (p=0.037) compared to untreated controls measured by the expression of either GATA-3 for Th2 or the RORγt transcription factor for Th17 cells among effector-memory CD4 cells. The Fra-2 mice had lower levels of Th1 cells that express the transcription factor, Tbet, or Treg cells that express the transcription factor, Foxp3, compared to healthy controls. Expression of Tbet or Foxp3 did not change in a statistically significant manner by treatment with SQL (Data not shown). Other markers on splenocytes for which we could not prove a statistically significant effect from treatment on cells or surface markers which included total CD4, total CD8, or PD1 (which trended lower), CD69, and effector-memory or central memory cells among either CD4 or CD8 cell T cell subsets (Data not shown).

### Soquelitinib is effective in treating imiquimod-induced psoriasiform skin lesions

We investigated the effectiveness of SQL to treat psoriasis-like skin inflammation using an imiquimod (IMQ)-induced murine model of psoriasis, in which IMQ, a TLR7 agonist^22^, induces IL-17 expression and Th17 cell differentiation in the skin lesions^23^. Human psoriasis is also IL-17 dependent^24^. We applied imiquimod cream on mouse back skin and compared the clinical and histopathological features of different treatment groups. Either the intermediate dose of SQL-formulated chow or control chow was given 6 days before applying the topical IMQ (see Methods). As shown in Figure 3A, modified clinical PASI scores for disease severity showed that SQL treatment significantly reduced skin thickness induced by IMQ. As expected, dexamethasone, a positive control, showed the greatest improvement of disease severity. Consistent with reduced clinical score, quantification of dermal inflammation, epidermal thickness, epidermal erosion, and signs of keratinocyte activation such as epidermal hyperplasia or epidermal hyperkeratosis revealed significant histological disease reduction within skin lesions from the SQL and the dexamethasone-treated groups (Figures 3B and 3C). Moreover, we found that SQL treatment significantly inhibited extension of epidermal inflammation into the underlying superficial dermis (Figure 3C). Because IMQ is also a potent immune activator, application of IMQ led to splenomegaly, a sign of enhanced systemic immunoreactions as shown in supplemental Figure 3. Interestingly, SQL treatment also reduced the systemic effect, leading to a decrease in spleen weight which did not achieve statistical significance (Supplemental Figure 3). Collectively, we demonstrated that pharmacological inhibition of ITK activity using SQL can decrease IMQ-induced psoriasis-like skin lesion indicators such as skin thickening and epidermal hyperplasia.

### SQL treatment inhibits Th17 cell differentiation

Evaluation of the IMQ-induced psoriasis model suggested that SQL is efficacious in Th17 cell-mediated disease. We further characterized the effect of SQL on Th17 cell differentiation *in vitro*. As shown in Figure 4D, using flow cytometry, we examined the expression of RORγt protein, a master transcription factor responsible for developing Th17 cells. We sorted naïve (CD44^low^CD62L^hi^) CD4+ T cells isolated from wild-type mice that were stimulated and cultured in Th17 polarizing conditions. RORγt protein expression was markedly reduced in SQL-treated Th17 cells under the same Th17-inducing cell culture conditions. Furthermore, we found SQL inhibited IL-17A cytokine produced in the culture supernatant in a dose dependent manner as the levels of IL-17A were virtually undetectable in cells cultured in the presence of 5 μM SQL (Figure 4E). In line with defective Th17 differentiation following SQL treatment, surface expression of CCR6, a chemokine receptor that plays a role in Th17 migration^25^, was greatly reduced in SQL-treated Th17 cells (Figure 4F). Collectively, our findings showed that SQL treatment regulates expression of RORγt, CCR6, and IL-17 production in Th17 cells.

**Figure 4.**
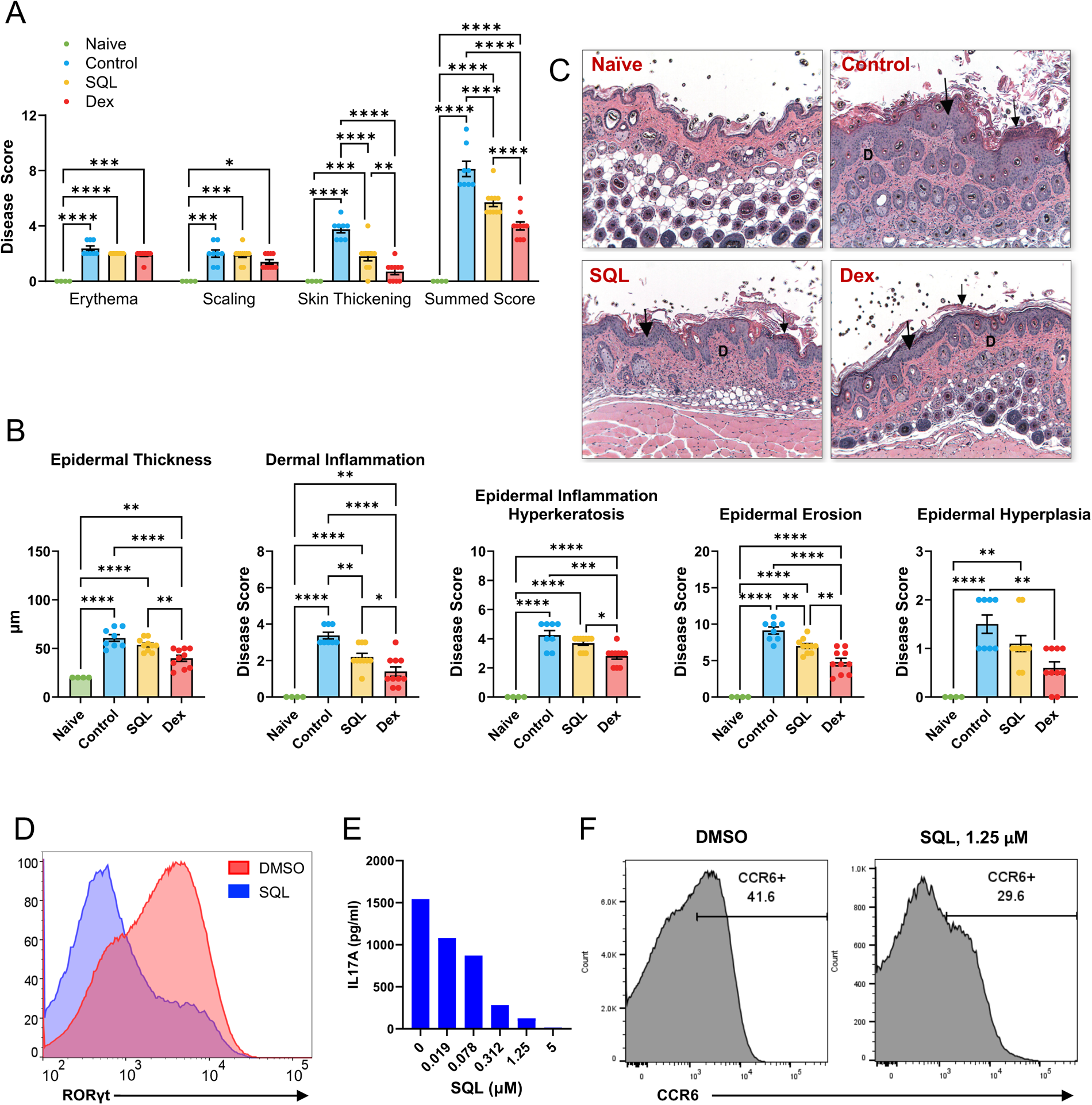
Soquelitinib decreases psoriasis-like skin inflammation induced by imiquimod. (A) Soquelitinib reduces overall modified PASI score. Mice skin Psoriasis Area and Severity Index (PASI) scores are based on erythema, scaling, and skin thickening and summed scores are shown. n=4–10. (B) Soquelitinib reduces dermal inflammation. Histological analysis of several parameters such as epidermal thickness, epidermal inflammation hyperkeratosis, epidermal hyperplasia, and dermal inflammation is shown. (C) Representative H&E staining of lesion skin sections is shown. Epidermal inflammation hyperkeratosis (small arrow), epidermal hyperplasia (large arrow) and dermal inflammation are shown. Selective ITK inhibition by soquelitinib impairs Th17 cell differentiation. (D) Histogram overlay of RORγt expression from wild-type naïve CD4 T cells differentiated under Th17 conditions in the presence of either DMSO or 1.25 μM of soquelitinib (SQL) for 6 days is shown. (E) IL-17A production in supernatants collected from Th17 cells treated with either DMSO or varying concentrations of soquelitinib (SQL) is shown. (F) Flow analysis of CCR6 expression from Th17 cells treated with either DMSO or 1.25 μM of soquelitinib (SQL) for 6 days is shown. Data are representative of two independent experiments. Statistical significance was determined by one-way ANOVA. Each dot in a bar graph represents one mouse. D=dermis. * p<0.05; ** p<0.01; *** p<0.001, **** p<0.0001.

### Soquelitinib reduces acute graft-versus-host disease

We next tested SQL in a graft-versus-host disease (GVHD) model. To investigate this, we employed the widely recognized MHC-disparate GVHD models using C57BL/6 (B6, H-2Kb) mice as donors and BALB/c (H-2Kd) mice as recipients. The recipient mice were treated with an intermediate dose of SQL-formulated chow or control chow starting from day -7 until day 90 following allogeneic hematopoietic cell transplantation (allo-HCT). We observed a significant improvement in survival rates (Figure 5A) accompanied by a corresponding decrease in clinical GVHD score (Figure 5B) in mice treated with SQL compared to the control group. The recipients treated with SQL exhibited reduced GVHD-related pathology compared to the control recipients on day 7 post allo-HCT (Figure 5C). The SQL group showed lower GVHD scores due to the attenuation of liver, small intestine, and large intestine disease, while the skin did not show significant amelioration (Figure 5C). T cell infiltration, as measured by CD3 staining, appeared to be reduced in SQL recipients (Figure 5D). The T cell infiltration is quantified in Figure 5E for the large or small intestine. To validate these findings, we utilized the B6 (H2Kb) into 129S1/SvImJ (129S1, H2Kb) GVHD model, which is MHC-matched but disparate for minor histocompatibility antigens^26^. Once again, recipients treated with SQL demonstrated improved survival rates in comparison to the control animals (Figure 5F). Notably, while there was one death in the SQL treated bone marrow group, no other mice experienced graft failure, indicating that SQL treatment does not impact engraftment, which is crucial when used as an prophylactic agent in the context of allo-HCT. To assess the potential impact of SQL on the graft-versus-tumor (GVT) effect, we conducted a study using a graft-versus-lymphoma model with a BALB/c-derived B-cell lymphoma cell line, A20. We observed long-term control of lymphoma exclusively in mice that received T cells (Figure 5G). Additionally, when SQL was administered to recipients receiving T cells, they demonstrated anti-tumor effect similar to the control group, but with reduced GVHD, resulting in improved survival compared to the control group (Figure 5F). These findings suggest that SQL does not significantly impair the GVT response. Overall, these results suggest that SQL effectively alleviates GVHD without compromising engraftment and GVT effect.

**Figure 5.**
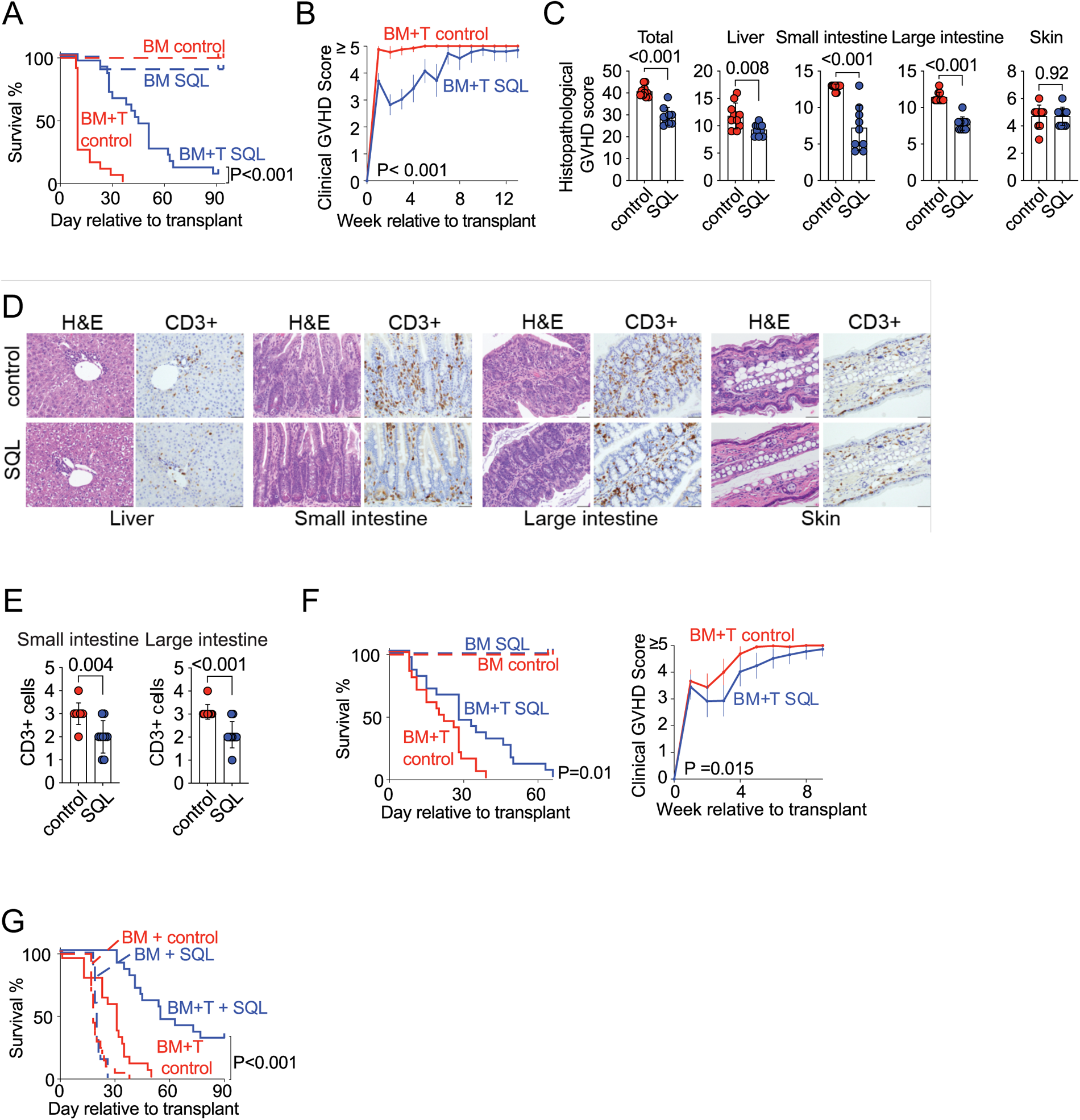
Soquelitinib treatment reduces GVHD severity. Sub**-**lethally irradiated BALB/c mice received B6 BM alone (BM which has been T cell depleted) or in combination with T cells (BM+T). (A) Survival and (B) clinical GVHD scores of BALBc mice that were treated with either SQL formulated or control diet from day -7 to day 90 relative to transplant. (C) Organ-specific and compound histopathological scores at day 7 post-transplant. (D) Representative histology images of transplanted mice. HE stain and CD3 stain. (E) CD3 positive T cells in the large or small intestine of SQL-treated mice or controls. (F) Survival and clinical GVHD scores of 129 mice receiving B6 marrow. (G) Sub-lethally irradiated BALB/c mice received B6 BM alone (BM) or in combination with T cells (BM+T) + A20 lymphoma cells. Each dot in a bar graph represents an individual mouse.

Subsequently, we investigated the effect of SQL treatment on alloreactive T cells, with a specific focus on T cell proliferation and activation. In the context of GVHD, there is an initial proliferation of donor CD4+ and CD8+ T cells in the spleen, followed by their migration to the intestines, liver, and skin. Analyzing carboxyfluorescein (CFSE)-labeled T cells in the spleen three days post-transplant, we observed reduced proliferation of both CD4 and CD8 T cells (Figure 6A), as indicated by CFSE dilution, accompanied by decreased T cell activation (Figure 6B), as suggested by reduced CD25 expression. By day 7, we noted decreased T cell proliferation (Figure 6C), measured by Ki67, and reduced T cell activation (Figure 6D), measured by CD25, particularly in the small and large intestines. Furthermore, using a mixed lymphocyte reaction assay, we found that 10μM SQL reduced the proliferation of T cells exposed to lipopolysaccharide (LPS)-activated allogeneic dendritic cells (DCs) compared with the vehicle control, as determined by CFSE dilution (Figure 6E). This reduction was accompanied by decreased activation, as indicated by CD25 expression. Similar effects were observed when T cells were stimulated with CD3/CD28 beads (Figure 6F), suggesting a T cell intrinsic effect at high concentrations of SQL. Overall, these findings indicate that SQL reduces alloreactive T cell proliferation and activation.

**Figure 6.**
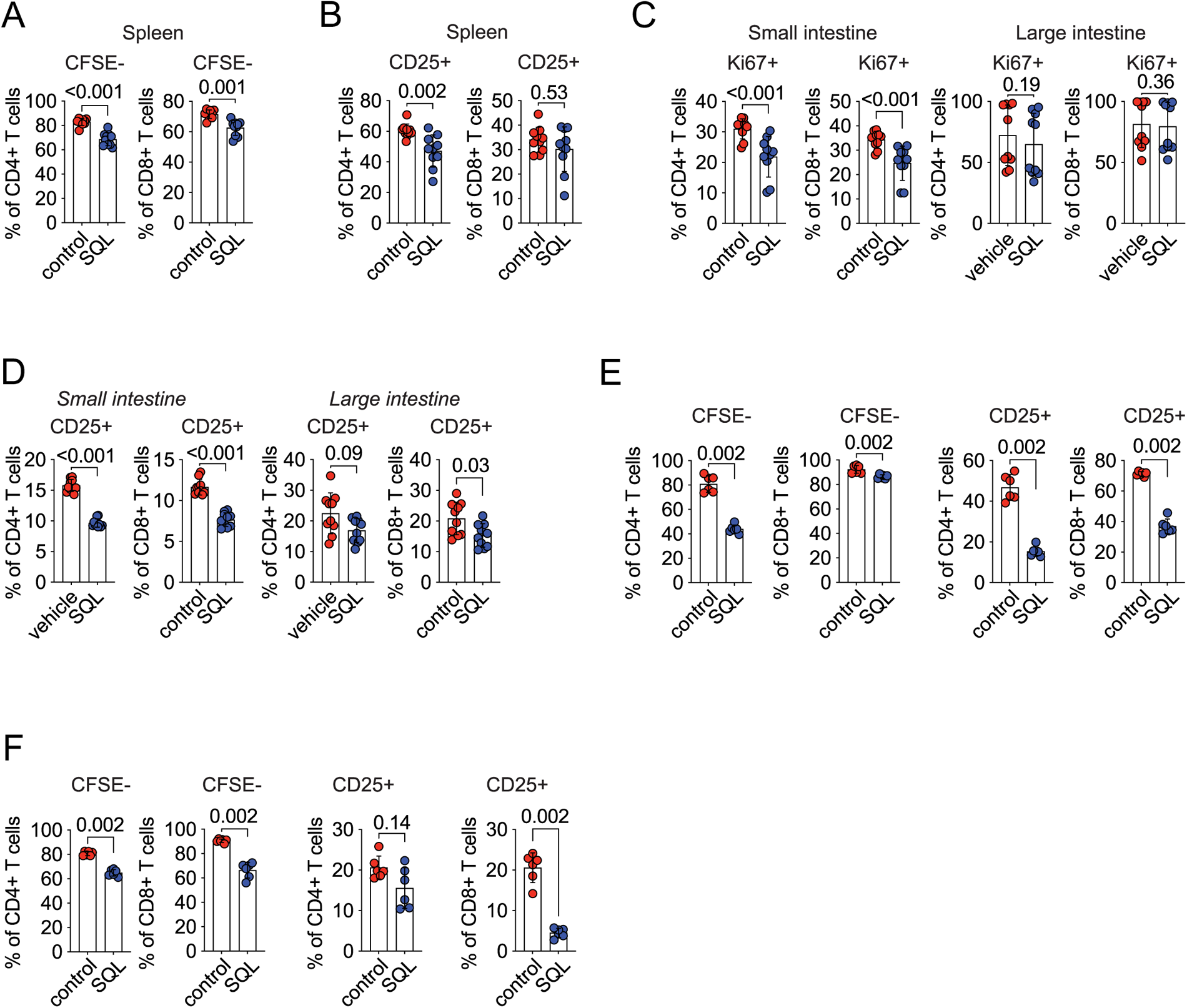
Soquelitinib reduces T cell activation and proliferation in GVHD. (A) Based on CFSE staining, SQL treatment reduces the number of splenic CD4 or CD8 cells that proliferate. (B) CD25, a marker of T cell activation, is significantly reduced on T cells in the spleen. The reduction of CD25 on CD8 T cells does not reach statistical significance. (C) Using Ki67 as a proliferation marker, SQL reduces the proliferation of either CD4 or CD8 T cells in the small intestine. Similar quantification of Ki67 is not statistically significantly different in the large intestine. (D) SQL reduces CD25 expression on either CD4 or CD8 cells in both the large and small intestine, although the measurements in the large intestine are of borderline significance (p=0.09). (E) MLR was performed using LPS activated DCs from BALB/c mice and T cells from B6 mice. Based on CFSE staining, SQL reduced the proliferation of either CD4 or CD8 T cells. Based on CD25 expression, SQL reduced the activation of either CD4 or CD8 lymphocytes. (F) Studies are similar to those depicted in Figure 6E, but the T cells have been activated by anti-CD3/anti-CD28 dynabeads. CFSE staining indicates reduced proliferation of CD4 or CD8 cells in the presence of SQL. CD25 staining indicates reduced activation of CD4 or CD8 lymphocytes in the presence of SQL, but the reduction in CD4 cells did not reach statistical significance (p=0.14). Each dot in a bar graph represents an individual mouse.

### DISCUSSION

ITK plays an important role in regulating signaling pathways downstream of TCR and in the differentiation of T helper (Th) cells^1,4,5^. Included in these Th cells are Th1 cells that express transcription factor T-bet and produce inflammatory cytokines such as IFNγ and Th2 cells that express transcription factor GATA-3 and secrete cytokines such as IL-4, IL-5 and IL-13 as well as Th17 cells that express transcription factor RORγt that secrete IL-17. In this capacity ITK plays a key role in fine-tuning T helper cell differentiation. ITK has also been shown to play a role in CD8 and ILC-2 function.

Our studies demonstrate the efficacy of an ITK inhibitor, SQL, in multiple murine models of T cell-mediated inflammatory disease including those that resemble acute and chronic asthma, pulmonary fibrosis, systemic sclerosis, psoriasis, and acute graft-versus-host disease. In additional studies beyond the scope of this report, we have noted efficacy of SQL in spontaneous atopic dermatitis in dogs (abstract)^27^, in the MRL/lpr murine model of systemic lupus erythematosus (abstract)^28^, and in a T-cell mediated model of colitis in mice. We believe that the mechanism that accounts for this benefit is multifactorial and probably varies for each model. This mechanism includes a reduction in the production of Th2 (see Figures 1B, 1E, 2F, and 3G) and Th17 cytokines (see Figures 3G, 4D, 4E, and 4F). An attractive feature of SQL is relative preservation of Th1 production at lower dosages of SQL^9,29,30^ which should reduce the potential for infectious complications due to immunosuppression. Indeed, in an ongoing clinical trial with SQL for the treatment of relapsed, refractory T cell lymphoma, we have demonstrated anti-tumor activity without an increased incidence of opportunistic infection (NCT03952078)^30^. The development of selective and effective inhibitors targeting specific kinases such as BTK, JAKs, and EGFR has revolutionized the treatment of numerous diseases^31^. In the aGVHD model, SQL reduces the activation and proliferation of T cells^9^ (see also Figure 6A). In asthma models, SQL inhibits the production of inflammatory cytokines such as IL-6 and TNFα (Figure 1B and 1E). These results suggest that inhibiting ITK could be a promising therapeutic approach for a wide range of T-cell-mediated disorders.

One of the key features of SQL is its selectivity towards ITK. While other inhibitors targeting related kinases have been developed, SQL demonstrates selectivity for ITK compared to these other kinase inhibitors targeting the TEC family including BTK, EGFR, and JAKs^9^. Members of the TEC family contain a cysteine residue in similar positions in the ATP binding site. SQL forms a covalent bond with the cysteine residue only in ITK accounting for its potency and selectivity. This selectivity is crucial in minimizing off-target effects and potential toxicity. At least one prior study evaluating ITK inhibition as a treatment for asthma in mice failed to show efficacy^32^. The lack of specificity of the studied inhibitor might have contributed to its lack of benefit in this study. The critical role of ITK in affecting the expression of the transcription factor, GATA-3, has been well established by us and others^9^. Thus, SQL has potential benefit in treating Th2 or innate lymphoid cell-2 mediated diseases which include asthma^33^, atopic dermatitis^34^, eosinophilic esophagitis^34^, nasal polyposis^35^, prurigo nodularis, and chronic obstructive pulmonary disease with eosinophilia^36^. Each of these is improved by treatment with dupilumab, a monoclonal antibody that blocks the receptor for two of the Th2-derived cytokines, IL-4 and IL-13. Therefore, a drug that reduces all Th2 cytokines has the potential to also be effective for these indications.

In this study, we demonstrated SQL is efficacious in IMQ-induced skin inflammation. IL-17 inhibition is effective treatment for psoriasis^12^, ankylosing spondylitis^37^, and psoriatic arthritis. The inhibition of RORγt and the reduction of Th17 cytokines suggest that SQL has potential to treat these diseases as well.

Fibrosis is characteristic of multiple diseases affecting tissues including liver, lung, kidney, brain, skin, blood vessels and heart. Fibrosis is a complex process in which Th2-derived cytokines have been strongly implicated^10^. Our report shows that SQL is effective in two well characterized models of fibrosis, pulmonary fibrosis secondary to bleomycin and peri-arterial and lung fibrosis in mice which over-express the transcription factor, Fra-2.

Despite the use of prophylactic immunosuppression, GVHD remains a significant challenge in allogeneic hematopoietic cell transplantation (allo-HCT), which is a potentially curative treatment for various hematologic malignancies and disorders. In acute GVHD, donor T cells attack host tissue and affect multiple organs, particularly the skin, liver, and gastrointestinal tract. This complication poses a substantial impediment to the success of allo-HCT, highlighting the need for effective strategies to mitigate GVHD without increasing the risk of infection and relapse, thus improving patient outcomes. Recent studies using ITK-/-in T cell graft or short-term exposure of donor graft to an ITK inhibitor have suggested that targeting ITK may be a promising approach for preventing GVHD^38–40^. Indeed, we demonstrated in a murine aGVHD model that treatment with SQL can decrease alloreactive T cell activation and proliferation, consequently reducing the incidence of acute GVHD. We do not believe that SQL acts directly on T cells to limit proliferation or viability except at very high concentrations. Rather, the reduced T cell proliferation observed in the GVHD model is relative to T cells that are being activated by cytokines and other inflammatory mediators.

A possible limitation of these studies is that the disease models were assessed at several different laboratories based on the expertise of each site. While this could be regarded as a drawback, we view it as a strength which indicates the reproducible benefit of SQL. In most of the models, we elected to deliver SQL within the chow as we have carefully assessed daily food consumption and titrated the dosage accordingly in order to achieve desired plasma concentrations. Dosing by gavage was also effective as in the pulmonary fibrosis model. In several of the studies reported above, the higher dose of SQL was less effective than the lower dosage. We have also observed this previously^9^. SQL preferentially inhibits Th2 cytokine production, but higher doses can also block the production of Th1 cytokines^9^. Thus, the highest tolerated dosage is not necessarily the most efficacious dosage.

In summary, the development of the highly selective and effective ITK inhibitor, soquelitinib, represents a promising new approach for the treatment of T cell-mediated diseases. The covalent binding mechanism and high occupancy of the inhibitor^9^, combined with its favorable safety profile, suggest that it may have potential as a targeted therapy for a range of diseases in which dysregulated T cell activation plays a role. Further studies are needed to fully evaluate its clinical efficacy and safety.

## METHODS

### Mice

For GVHD experiments, B6, BALBc, and 129S1 mouse strains were obtained from the Jackson Laboratory. The psoriasis and IPF models were studied in female C57BL/6 mice (Charles River) while the asthma models were studied in female BALB/C mice (Charles River). The Fra-2 mice were bred and housed at INSERM. Only female mice aged 6-8 weeks were studied for all models with the exception of the Fra-2 model for which treatment began at age 11 weeks. All animal studies were reviewed and approved by the locally appropriate review panel.

### Soquelitinib treatment and formulations

SQL was formulated as a solution for oral gavage in water containing tween, propylene glycol and cellulose, and administered to mice at doses of either 10mg/kg or 30mg/kg. Pharmacokinetic studies showed that these doses produced plasma Cmax levels of SQL of 600ng/ml and 3500ng/ml, respectively (median, 5 animals/group). Cmax levels were used because SQL is a covalent drug that irreversibly binds to the target.

For chronic dosing regimens, a mouse chow formulation was developed. Mice ingested chow containing SQL continuously during the period of dosing. Chow formulations of 1.2g SQL/kg chow (low dose); 2.4g SQL/kg chow (intermediate dose) and 7.2g SQL/kg chow (high dose) were evaluated. These formulations produced plasma Cmax levels of 2000ng/ml, 3800ng/ml (median, 5 animals/group) and approximately 7600ng/ml, respectively. Levels in the mice receiving the high dose were estimated based on food consumption, which was reduced in this group.

### OVA-induced asthma model

The protocol was based on a report previously described^15^. For acute asthma model, wild-type BALB/c female mice were randomly divided into 5 groups (Group A-E, Figure 1A) and were sensitized intraperitoneally twice at 5-day intervals with either PBS or 10 μg of Ovalbumin (OVA, InvivoGen) together with 1 mg of aluminum hydroxide adjuvant (InvivoGen) on day 0 and day 5. On day 6, mice were treated by oral gavage with either vehicle alone, or one of two different doses of SQL (10 mg/kg or 30 mg/kg) in solution. Doses were administered once daily for ten consecutive days in a volume of 200μl. From day 12 to 15, the sensitized mice were challenged intranasally with either PBS or OVA solution in PBS (40 μg of OVA). As a treatment positive control, group E mice were treated with dexamethasone (1 mg/kg, InvivoGen) intraperitoneally once daily from day 12 to 15. On day 16 (24 hours after the last OVA challenge), mice were euthanized and BALF and lungs were collected for cytokine production, and histopathology.

For the chronic asthma model, the study design was outlined in Figure 1E. In brief, wild-type BALB/c female mice were sensitized intraperitoneally twice at 7-day intervals with either PBS (Group A) or 40 μg of OVA (Group B-F) together with 1 mg of aluminum hydroxide adjuvant (InvivoGen) on day 0 and day 7. Treatment began following ad libitum access to either control chow containing vehicle control (Group A and B), or medicated chow containing one of three different doses (low, intermediate, or high dose) of SQL (Group C-E), for 45 days until study was terminated on day 53. From day 15 to day 52, the sensitized mice were challenged intranasally with either PBS or OVA solution in PBS (40 μg of OVA) twice a week. Similarly, on day 53, mice were euthanized and BALF, and lungs were collected for cytokine production, and histopathology.

### Bleomycin-induced pulmonary fibrosis (IPF) model

Forty-five eight-week-old C57BL6 female mice were randomly divided into 5 groups (Group A-E, Figure 2A). On day 0, 40 animals were administered bleomycin (1.5U/kg, Hospira) via the oropharyngeal route and five animals were administered normal saline via the oropharyngeal route as Group 1. On day 6, bleomycin administered animals were distributed into Groups B-E based on an equal distribution of animal mean body weight. Treatment with either vehicle control, one of two different doses of SQL (either 10 mg/kg or 30 mg/kg) in solution or nintedanib (50 mg/kg, Axon Medchem) was begun on day 7 post bleomycin administration once daily (QD) via oral gavage and continued until the study was terminated on day 21 as described in Figure 2A. On day 21, all surviving animals were harvested within 1 to 3 hours post final dosing. Animals were then sacrificed and lungs (except the post caval lobe) were weighed and used for histopathology. BALF was also collected and centrifuged for differential cell counts. Differential leukocyte number was determined by cytospin of cells recovered from BALF of individual mice. Post caval lobes were instantly frozen in liquid nitrogen for subsequent RT-PCR mRNA quantification.

### Systemic sclerosis model

Mice that over-express the transcription factor, Fra-2, and develop vascular injury followed by fibrosis have been previously described^19,20^. Antibodies used for studies related to the Fra-2 model were purchased from Biolegend. Assessment of fibrosing alveolitis by Micro-CT images were obtained with a Perkin Elmer Quantum FX system (Caliper Life Sciences, GmbH, Mainz, Germany). The animals were placed in the supine position on the CT table and sedated with 3– 4% isofurane anesthesia under 0.5–1.5 L/min for induction by a nose cone. Anesthesia was maintained with 2.5–3% isofurane under 400–800 mL/min during the acquisition. Images were acquired with the following parameters: 90 kV X-ray source voltage, 160 μA current. Means of lung density were achieved by evaluation of all CT scans acquired from the apices to the bases of the lungs. RVSP (right ventricular systolic pressure) and heart rate were measured in unventilated mice under isofurane anesthesia (1.5–2.5%, 2 L O2/min) using a closed chest technique. A catheter (1.4-F catheter; Millar Instruments Inc., Houston, TX) was introduced into the jugular vein and directed to the right ventricle. Morphometric analyses were performed on paraffin embedded lung sections stained with hematoxylin and eosin and alpha smooth muscle actin (α-SMA).

### Imiquimod-induced murine model of psoriasis

Female C56BL/6 mice were given access to either control chow containing vehicle control (Naïve, Control, medicated chow containing SQL at an intermediate dose of 2.4 gm SQL/kg of chow beginning on day -6. These chow diets were available to the animals at all times (ad libitum) until study termination on day 8. On day -1, animals in dexamethasone group were treated with dexamethasone (1 mg/kg) daily (QD) by oral gavage. On day 0, animals were given baseline modified Psoriasis Area and Severity Index (PASI) scores based on three clinical signs (erythema, scaling, and skin thickening), and calipered for baseline measurements. Additionally, on day 0, mice from Dexamethasone group were begun on dexamethasone daily by oral gavage (1 mg/kg, InvivoGen) as a positive control. On days 1–7, the majority of animals except mice in the Naïve Group had 5% Imiquimod (IMQ) cream (Perrigo) applied to the hair-free skin of the back and rubbed in until absorbed. PASI scores and caliper measurements were recorded for skin thickness on days 0, 2, 4, 6, and 8. Following final PASI scores and caliper measurements on day 8, the mice were euthanized, and skin samples were collected for histopathology. The modified PASI scoring system is used to evaluate disease severity. Parameters of erythema, scaling and skin thickening were graded from 0 to 4, where: 0 = none, 1 = mild, 2 = moderate, 3 = severe in each parameter. The cumulative score indicates the severity of disease from factors of erythema, scaling and skin thickening (total = 0–12).

### GVHD and GVT model

The GVHD experiments at MSKCC were performed as previously described^41^. The conditioning regimens involved split-dosed sub-lethal irradiation, with 900 cGy administered to BALB/c recipients and 1000 cGy to 129S1 recipients. Bone marrow (BM) cells were depleted of T cells using anti-Thy-1.2 and Low-Tox-M rabbit complement (CEDARLANE Laboratories). Donor T cells were isolated from splenocytes using the Pan T Cell Isolation Kit II (Miltenyi), resulting in T cell purity averaging >90%. BALBc hosts received 10 x 10^6^ BM cells with or without 1 x 10^6^ T cells, while 129S1 hosts received 5 x 10^6^ BM cells with or without 4 x 10^6^ T cells via tail vein injection. Mice were monitored daily for survival and assessed weekly for GVHD clinical scores, as previously reported^41^. Recipients were provided with either control chow containing vehicle control or medicated chow containing the intermediate dose of SQL from day -7 to day 90 relative to BM+T or until the day of harvest. In the GVT experiments, mice were administered BM alone or BM with T cells via retroorbital injection, and they received an additional injection of 5 x 10^6^ A20 lymphoma cells via the tail vein on day 0. In the T cell proliferation experiment, donor T cells were labeled with 5 μM Cell Trace CFSE (Invitrogen) according to the manufacturer’s instructions. Additionally, the T cell dose was increased to 5 x 10^6^ T cells.

### GVHD histopathology

Approximately 2 cm segments of the distal small intestine and proximal colon, along with samples of the liver and ear, were collected on day 7 after allo-HCT. The harvested tissues were fixed in 10% formalin, embedded in paraffin, and sectioned at a thickness of 5 μm. Hematoxylin and eosin (H&E) staining, Ki67 and CD3 immunohistochemical (IHC) staining and terminal deoxyribonucleotidyl transferase mediated dUTP-biotin nick end labeling (TUNEL) assays were performed by the Laboratory of Comparative Pathology facility at MSKCC, following previously described methods^41^. TUNEL staining was a component of the scoring for histopathology. The slides were evaluated by a board-certified veterinary pathologist who was blinded to the group treatments during the assessment.

### MLR and T cell activation assay for GVHD studies

*Preparation of bone marrow-derived dendritic cells (BMDCs):* BMDCs were prepared from WT-BALB/c mice. BM cells were isolated and cultured in DMEM medium supplemented with 20ng/ml recombinant GM-CSF (Peptrotech) and 5ng/ml IL-4 (Peptrotech) for 6 days. On the 6th day, BMDCs were matured using 50ng/ml of LPS (Sigma) for 24 hours. *Isolation and Activation of T Cells:* T cells were isolated from the spleen of B6 mice using the Pan T cell isolation kit (Miltenyi), following the manufacturer’s protocol. The isolated T cells (2 × 10^5^) were either cocultured with irradiated (20Gy) BALB/c mice BMDCs (5 × 10^4^) or activated using mouse T-activator CD3/CD28 Dynabeads (Gibco) at a 1-to-1 bead-to-cell ratio. *Assessment of T Cell Proliferation:* T cells were labeled with 5 μM Cell Trace CFSE (Invitrogen) as per the manufacturer’s instructions. On day 4 of the culture, the cells were transferred into V-bottom plates (Fisher), pelleted by centrifugation, and incubated with an antibody staining mix containing DAPI (ThermoFisher) viability dye diluted in PBS for 20 minutes at 4 °C.

### In Vitro Th17 differentiation

For Th17 cell differentiation, CD4+CD62L^high^CD44^low/-^ T cells were purified from the spleen of wild-type BALB/c mice post staining with antibodies to mouse CD3 (17A2, Biolegend), CD4 (RM4-5, Biolegend), CD44 (IM7, Biolegend), CD62L (MEL-14, Biolegend), followed by sorting of cells using the SONY SH800 sorter. 3x10^5^ sorted naïve CD4 T cells were stimulated with plate-bound anti-CD3 (145-2C11, Cytek) (5 μg/ml) and soluble anti-CD28 (clone 37.51, Biolegend) (1 μg/ml) in the presence of Th17 polarizing cocktails containing recombinant mouse IL-6 (10 ng/ml, Biolegend), mouse IL-1β (10 ng/ml, Biolegend), mouse IL-23 (10 ng/ml, Biolegend), human TGFβ-1 (5 ng/ml, Miltenyi Biotec), anti-mouse IL-4 (11B11, Biolegend) and anti-mouse IFNγ (XMG1.2, Biolegend) neutralizing antibodies (10 μg/ml each, Biolegend) with or without SQL for 6 days. Culture supernatants were collected, and the concentrations of cytokines were assessed by MSD as described below.

### Flow cytometry

Cells harvested from *in vitro* Th17 differentiated cells were stained with the following antibodies against mouse CD3 (17A2, Biolegend), CD4 (GK1.5, Biolegend), CCR6 (29-2L17, Biolegend), RORγt (Q21-559, BD). Fixable Viability Dye eFuor780 (eBioscience) was used to exclude dead cells before analysis. Single cell suspension from spleen, mesenteric LN (MLN) and intestine tissues were performed as previously described^41^. Cells were stained with the following antibodies against mouse CD4, CD8, CD25. Ghost Dye™ Violet 510 (TonboBio) was used to exclude dead cells before analysis.

For intracellular staining of RORγt, cells were first stained with surface markers and then fix/permeabilized using the Foxp3/Transcription Factor Staining kit (eBioscience) according to the manufacturer’s protocol.

### Cytokine analyses

For mouse asthma studies, cytokine levels were measured by MSD (Meso Scale Diagnostics) platform using a custom designed mouse cytokine U plex kit and quantified by normalization with total protein levels in BALF. For the in vitro Th17 differentiation culture, levels of IL-17A in the supernatants were measured by MSD directly according to the manufacturer’s instructions.

### RNA isolation and quantitative PCR

Total RNA was extracted from snap-frozen post caval lobe using RNeasy kit (Qiagen) according to the manufacturer’s instructions. cDNA was synthesized with SuperScript III Reverse Transcriptase (Invitrogen). Then gene expression was determined by SYBR green real-time PCR using SYBR^TM^ Premix kit (BioRad) and was expressed using the 2 ^-ΔΔ^Ct method. The primer sequences used in this study were shown as following: beta actin (forward)-5’ggctgtattcccctccatcg 3’; beta actin (reverse)-5’ccagttggtaacaatgccatgt 3’; CD4 (forward)-5’ tcctagctgtcactcaaggga 3’; CD4 (reverse)-5’ tcagagaacttccaggtgaaga 3’; GATA-3 (forward)-5’ctcggccattcgtacatggaa 3’; GATA-3 (reverse)-5’ ggatacctctgcaccgtagc 3’; MMP2 (forward)-5’ caagttccccggcgatgtc 3’; MMP2 (reverse)-5’ ttctggtcaaggtcacctgtc 3’; TGFβ1 (forward)-5’ ctcccgtggcttctagtgc 3’; TGFβ1 (reverse)-5’ gccttagtttggacaggatctg 3’ All primers were purchased from IDT. Results from MMP2 and TGFb1 were normalized to beta-actin expression and presented as fold increase in mRNA expression relative to the level detected in control mice. Data from GATA-3 relative expression were further normalized to CD4 relative expression and presented as ratio of GATA-3 to CD4. Data were obtained from triplicate samples from 9–10 mice in each group.

### Histology for the IPF model

Lung, skin tissues were fixed in 10% (v/v) formalin for over 24h and embedded in paraffin. Sections were stained with hematoxylin and eosin (H&E) or Masson’s Trichrome (MT). For OVA-induced asthma model, lung inflammation was assessed by three parameters: peribronchial and perivascular inflammation and alveolitis. A score of 0–5 was given to each parameter. The maximum cumulative score of lung disease index is 15. For Bleomycin-induced fibrosis model, severity of fibrosis in lung tissue was assessed by evaluating the slides at 20X magnification and scoring 15 representative (random) fields for the average or most severe finding in the field using modified Ashcroft Score system^42^ with mean values ranging from 0 to 8. Fibrosis score was quantified by examining fibrotic connective tissue in the MT-stained sections according to the following scale: 0 = normal levels of collagen, 1 = <10% of the lung affected, 2 = 10–25% of the lung affected, 3 = 26–50% of the lung affected, 4 = 51–75% of the lung affected, 5 = >75% of the lung affected. For IMQ-induced psoriasis model, stained H&E sections from fixed skin pieces were used for pathology evaluation and scoring in five different parameters including epidermal thickness, epidermal hyperplasia, epidermal hyperkarotosis, epidermal erosion, and dermal inflammation. A 5-point scale was used where: 0 = normal, 1 = minimum (6–10% of the total surface area), 2 = mild (11–25% of the total surface area), 3 = moderate (26–50% of the total surface area), 4 = marked (51–75% of the total surface area), and 5 = severe (>75% of the total surface area) in each parameter.

### Statistics

The data are displayed as means ± SEM, unless otherwise indicated. All statistical analyses were performed using GraphPad Prism Software. Significance was determined using an unpaired two-tailed Student t-test, unless otherwise indicated. Differences were considered statistically significant at p<0.05. P values are reported as: ns. not significant, p> 0.1, *p <0.05, **p <0.01, ***p <0.001, ****p,0.0001.

