## Supplementary figures and images for "Soquelitinib, A Selective Inhibitor of Interleukin-2-Inducible T Cell Kinase (ITK), is Active in Several Murine Models of T Cell-Mediated Inflammatory Disease"

### Supplemental figures

#### Supplementary Figure 1

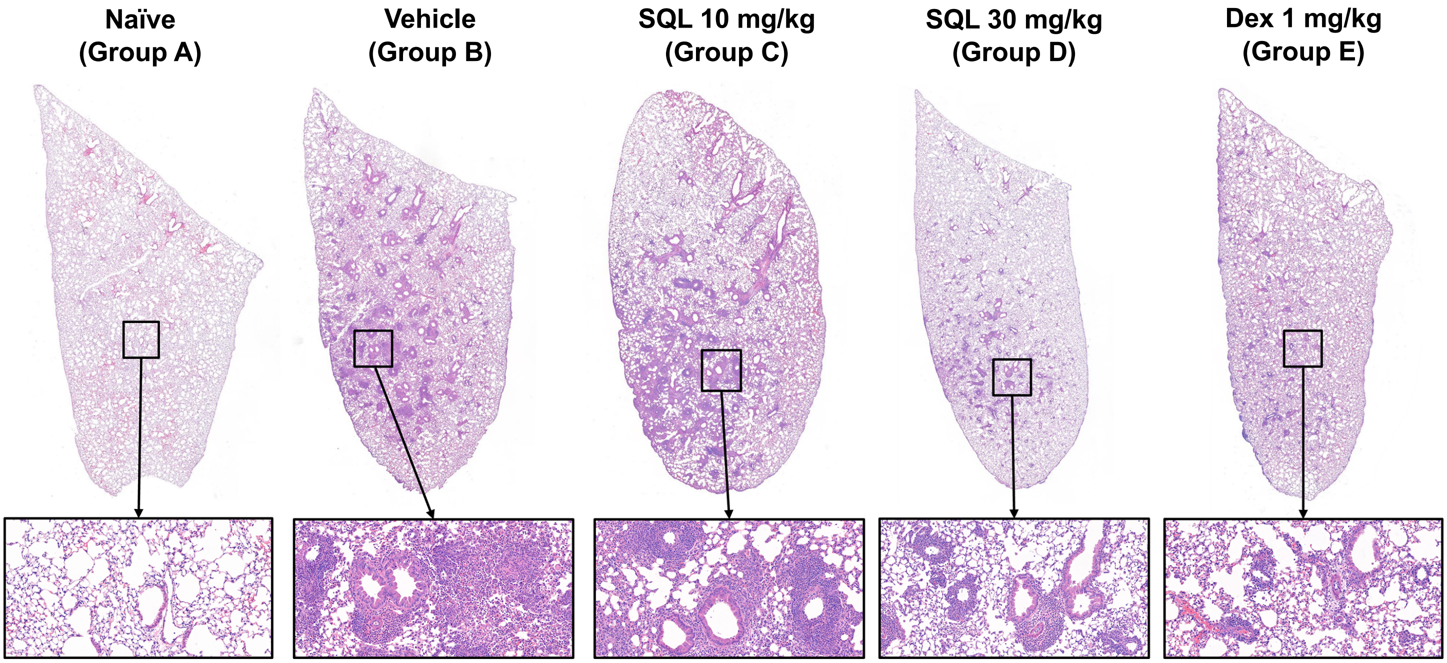

**Supplementary Figure 2**

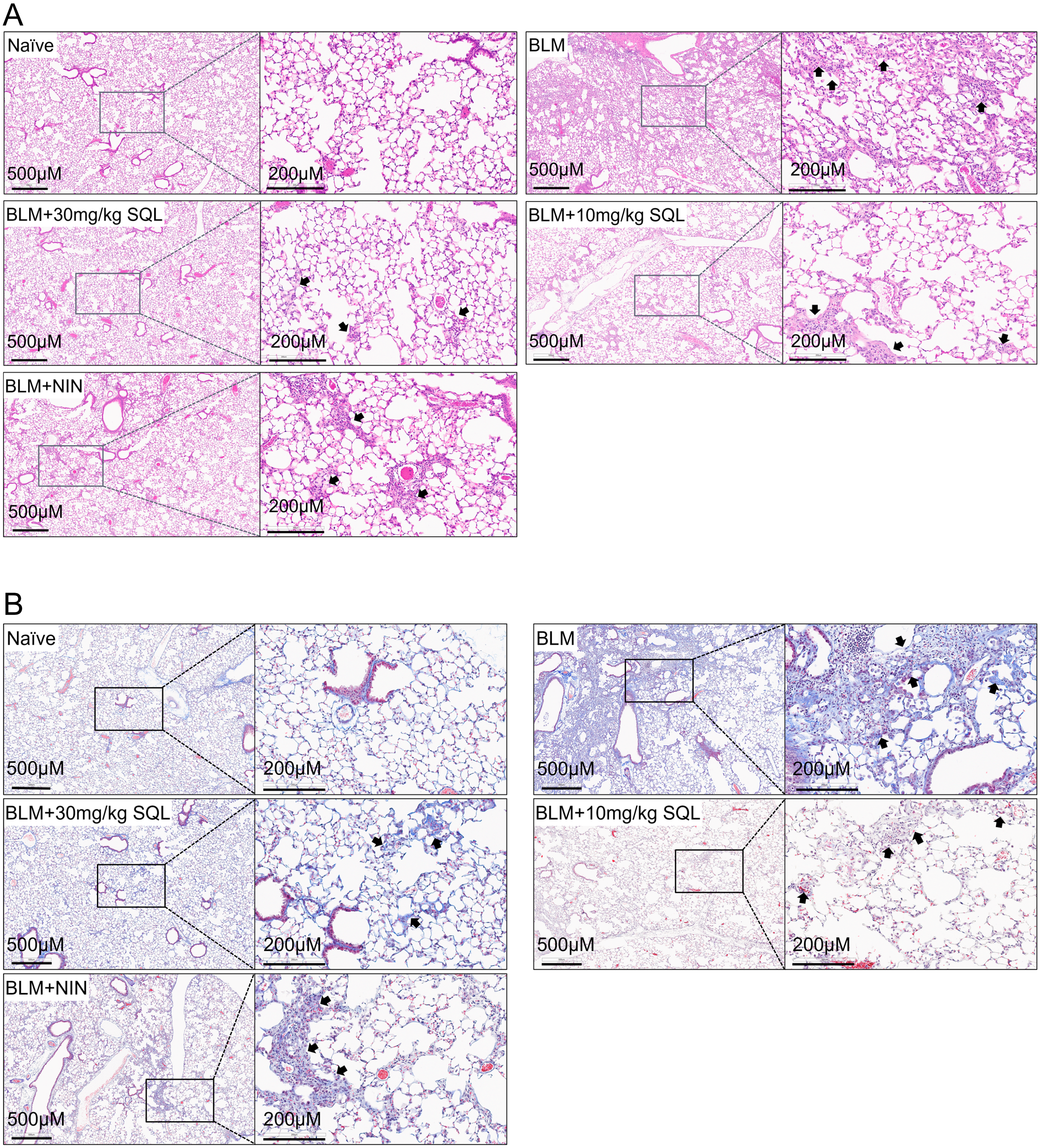

#### Supplementary Figure 3

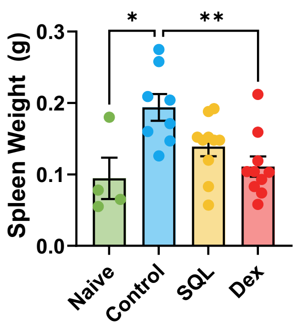
